# HiCAR: a robust and sensitive multi-omic co-assay for simultaneous measurement of transcriptome, chromatin accessibility, and cis-regulatory chromatin contacts

**DOI:** 10.1101/2020.11.02.366062

**Authors:** Xiaolin Wei, Yu Xiang, Armen Abnousi, Tongyu Sun, Xin Lin, Wei Li, Ming Hu, Yarui Diao

**Author notes:** These authors contributed equally to this work (Xiaolin Wei, Yu Xiang). Correspondence should be addressed to: M.H.; Y.D.

## Abstract

High-order chromatin organization plays a central role in regulating spatial-temporal gene expression by facilitating or constraining the interactions between cis-regulatory elements (cREs). cREs are often the accessible DNA sequences and can be identified at genome-wide scale with assays such as ATAC-seq, DHS-seq, and FAIRE-seq. However, it remains technically challenging to comprehensively identify the long-range interactions that occur between cREs, especially using low-input cells and in a cost effective manner. Here, we report HiCAR, <u>Hi</u>gh-throughput <u>C</u>hromosome conformation capture on <u>A</u>ccessible DNA with m<u>R</u>NA-seq co-assay, which enables simultaneous mapping of chromatin accessibility and cRE anchored chromatin contacts. Notably, using the same input material, HiCAR also yields high-quality transcriptome data that represents the functional outputs of chromatin accessibility and interaction. Unlike immunoprecipitation-based methods such as HiChIP, PLAC-seq, and ChIA-PET, HiCAR does not require target-specific antibodies and thus can capture cis-regulatory contacts anchored on the accessible DNA regions associated with multiple epigenetic modifications and transcription factor binding. We compared HiCAR to another technology designed to capture interactions between accessible chromatin regions, called Trac-looping, and found that HiCAR yielded much more informative long-range cis-reads at similar sequencing depth, and requires far fewer cells as input. We applied HiCAR to H1 human embryonic stem cells (hESC) and identified 46,792 open-chromatin anchored loops at 5Kb resolution. Interestingly, we found that the poised cREs form extensive and significant chromatin interactions comparable to the active cREs. We further showed that the spatial interactive activity of cREs do not correlate with their transcriptional activity, enhancer activity, and chromatin accessibility. Additionally, we identified 2,096 <u>s</u>uper <u>inte</u>ractive regulatory (SINTER) loci showing abnormally high levels of chromatin interactivity and associated with unique epigenetic features. In summary, HiCAR is a robust, sensitive, and cost effective multi-omics coassay that can be used to study chromatin structure and function as well as gene expression.

## Introduction

Cis-regulatory elements (cREs), such as enhancers, promoters, insulators and silencers, play a critical role in regulating spatial-temporal gene expression in development and diseases^1–3^. CREs are characterized by the presence of “open” or accessible chromatin that is depleted of packaging nucleosome particles to make way for the binding of Transcription Factors (TFs) and a variety of epigenetic remodelers. These accessible chromatin regions can be identified by Assay for Transposase-Accessible Chromatin using sequencing (ATAC-Seq)^4^, DNase-Seq^5^ and FAIRE-Seq^6^ (Formaldehyde-Assisted Isolation of Regulatory Elements). CREs also form dynamic high-order chromatin interactions to precisely control the expression of distal target genes. The development of chromosome conformation capture (3C)-based technologies has greatly improved our understanding of the principles of high-order chromatin organization, and revealed how dynamic chromatin looping affects gene expression in a cell type specific manner. Hi-C is widely used to detect higher-order chromatin features^7,8^, and can identify cis-regulatory interactions, but extremely deep sequencing (billion of reads) is required to to resolve long-range cis-regulatory interactions at kilobase resolution. To reduce the sequencing costs associated with mapping cis-regulatory interactions, alternative methods can be used such as ChIA-PET, HiChIP, PLAC-seq and Capture-C^9,10 11–14^ However, these methods rely on ChIP-grade antibody or pre-designed capture probes to enrich a subset of chromatin interactions associated with specific proteins, histone modifications, or targeted genome regions. More recently, Trac-looping and Ocean-C have been developed to analyze interactions among accessible chromatin regions, independent of ChIP antibodies or capture probes^15,16^. While these methods do not require target specific pulldowns, their potential remains limited by high cell number requirements and relatively low portions of informative long-range cis reads. Thus, it prevents their application in low-input materials that are often collected from clinical samples and primary tissues. Moreover, none of the methods described above enable simultaneous measurements of gene expression from the same sample, which represent key functional outputs of cis-regulatory interactions. Therefore, a robust, sensitive, and cost effective method is urgently needed for comprehensive analysis of DNA accessibility and cRE anchored chromatin contacts, as well as the transcriptome that represents their functional outputs using low input material.

Here, we introduce a robust and sensitive method, called HiCAR, which allows genome-wide profiling of long-range chromatin interactions anchored on open chromatin regions. HiCAR requires only ~100,000 cells as input, and leverages principles of *in situ* HiC and ATAC-seq method and avoids many potentially loss-prone steps, such as adapter ligation and biotin-pull down. With similar sequencing depth, HiCAR outperforms Trac-looping^15^. by generating ~17-fold more (18.3%. V.s 1.1%) informative long-range (>20kb) cis-paired-end tag (cis-PET), even when starting from 1000-fold fewer cells. As a multi-omics co-assay, HiCAR also yields high-quality transcriptome and chromatin accessibility data from the same low-input starting material. Applying HiCAR to H1 human embryonic stem cell (hESC), we revealed novel features of chromatin structure and function related to epigenome, gene regulation, and high-order chromatin organization.

## Results

### Principle of HiCAR

As a proof-of-principle, we set out to perform HiCAR on H1 hESCs, because of the many genomic datasets available for this cell line that could be used to benchmark our approach^2,17^. As shown in Fig 1A, ~100,000 crosslinked cells were treated with Tn5 transposase assembled with a engineered DNA adaptor containing a Mosaic End (ME) sequence for Tn5 recognition^18^ as well as a flanking sequence that can be ligated to the CviQI-digested DNA fragment when supplemented with a splint oligo. After transposition, restriction enzyme digestion was performed with the 4-base cutter CviQI, followed by in situ proximity ligation to ligate Tn5 adaptors to proximal genomic DNA. After in situ ligation, crosslinks were reversed, the DNA was purified, digested by another 4-base cutter NlaIII, and circularized by re-ligation. The circularized DNA was used for PCR amplification to make HiCAR DNA libraries for next-generation-sequencing (NGS) using a previous method described earlier^19^. The PCR primers used for library amplification will anneal to the ME sequence and splint oligo sequence, respectively, and the chimeric DNA fragment will be amplified with one end derived from the CviQI digested genomic DNA (Fig 1A, blue, captured by Read 1 of each paired-end sequence) and the other end derived from the Tn5-tagmented open chromatin sequence (Fig 1A, red, captured by Read 2 of each paired-end sequence). PolyA RNAs from the cytoplasm and nucleoplasm were also collected during the procedure as indicated in Fig 1A, and then subjected to RNA-seq preparation using a protocol modified from SMART-seq2^20^

**Figure. 1.**
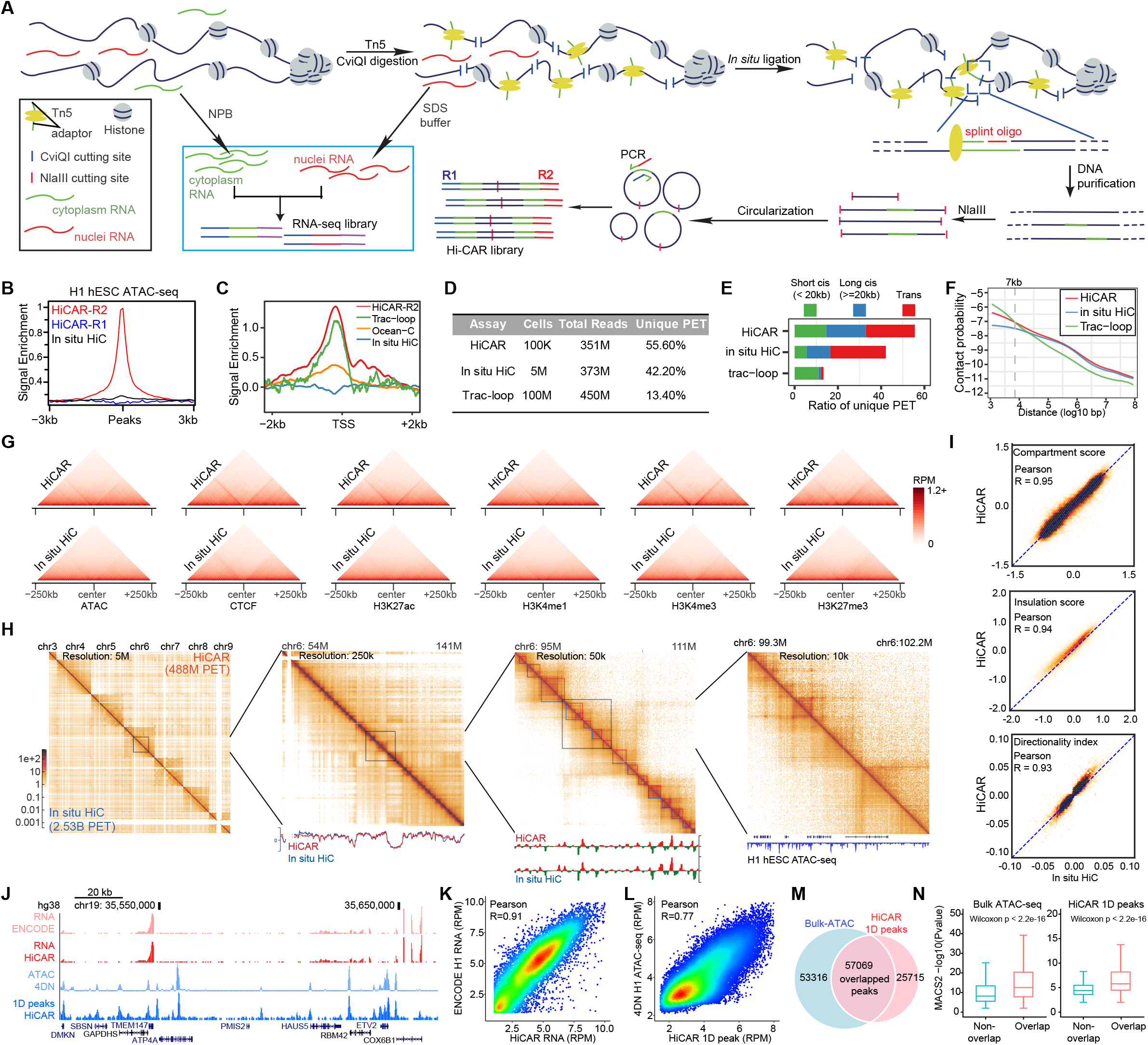
Schematic overview of HiCAR method. **(A)** In a HiCAR experiment, the crosslinked nuclei are treated by Tn5 transposase loaded with engineered DNA adapters, followed by restriction enzyme 4 base cutter CviQI digestion and in situ ligation. The Tn5 adapters can be ligated to the spatially proximal genomic DNA digested by CvQI. The ligated genomic DNA are purified after reverse crosslinking, and subjected to restriction enzyme digestion by another 4 base cutter NlaIII. Then resulting DNA fragments are circularized and PCR amplified. The DNA sequences amplified from the Tn5/ME sequence and splint oligo sequence are defined as R2 reads and R1 reads, respectively. The cytoplasm and nuclei RNA fractions are collected and pooled together for RNA-seq analysis **(B)** Normalized signal in a window of 3kb on each side of ATAC-seq peaks for HiCAR R2 (red), HiCAR R1 (blue) and *in situ* HiC (black). **(C)** Normalized signal in a window of 2kb around TSS for HiCAR R2 (Red), Trac-looping (green), Ocean-C (orange), *in situ* HiC (blue). **(D)** Comparison of input cells number and sequence outputs across different methods. **(E)** Percentage of unique short range cis, long range cis and trans reads ratio for HiCAR, *in situ* HiC and Trac-looping. **(F)** contact frequency as a function of distance measured by HiCAR, in suit HiC and Trac-looping. **(G)** Heatmap of aggregated contact matrix at a resolution of 10kb around indicated histone marker peakset. (top panel, HiCAR; bottom panel, *in situ* HiC). **(H)** Contact matrices of four successive zoom-in for H1. Each plot shows HiCAR data above the diagonal and *in situ* HiC data from 4DN data portal below the diagonal. The color represents normalized reads count. **(I)** Scatter plots show the global correlation of compartment scores (top panel), TAD insulation score (middle panel) and TAD directionality index (bottom panel) between HiCAR and *in situ* HiC. The r value: Pearson correlation coefficient. **(J)** A representative Genome Browser view of HiCAR RNA profile and ATAC profile in H1 cells. H1 bulk RNA-seq and ATAC data downloaded from ENCODE and 4DNA data portal are also shown. **(K-L)**. Scatter plots show the reads counts from HiCAR RNA profile and bulk polyA RNA-seq dataset **(K)** and HiCAR ATAC-profile and bulk ATAC-seq dataset **(L). (M)** Venn diagram showing the overlap of HiCAR ATAC peaks and bulk ATAC-seq peaks from 4DN data portal. **(N)** Boxplots show the MACS2 confidence scores of ATAC-seq peaks that overlap and do not overlap with each other.

In this study, we made three replicate HiCAR libraries and sequenced each of the HiCAR DNA libraries to a depth of ~350 million pair-end raw reads. We first examined the enrichment of reads around H1 open chromatin regions, analyzing the HiCAR read 1 (R1) and read 2 (R2), separately, as well as public in situ HiC data from H1 hESCs generated by 4DN^21^ as a negative control of no enrichment (Fig 1B). As expected, HiCAR R2 reads were highly enriched on the H1 hESC ATAC-seq peaks, while the R1 reads and in situ HiC show no enrichment at open chromatin regions (Figure 1B). This data confirmed that HiCAR successfully captured and enriched for interactions between open chromatin regions (R2) and other genomic regions (R1). We refer to these interactions below as “open-to-all” interactions. This is different from another method called Trac-looping^15^, which captures “open-to-open” interactions between pairs of open chromatin regions. Since open chromatin can be bound by multiple TFs and enriched for different histone modifications^22^, we asked if HiCAR could capture the chromatin interactions associated with distinct epigenome features. Indeed, we found that the HiCAR R2 reads, but not R1 reads, are highly enriched on H1 hESC H3K27ac, H3K3me1, H3K4me3, H3K27me3, RAD21, CTCF, NANOG and SOX2 ChIP-seq peaks (Fig S1A).

Next, we compared the efficiency of HiCAR to that of Trac-looping and Ocean-C, two methods that were published recently allowing for enrichment of long-range interactions anchored on open chromatin regions^15,16^. Because HiCAR, Trac-looping and Ocean-C were carried out in different cell lines, we decided to assess open chromatin enrichment efficiency by examining Transcription-start-site (TSS) signal enrichment, a metric which has been widely used as a quality control standard to compare signal-to-noise ratio of ATAC-seq across different cell types^23^. As shown in Fig 1C, both HiCAR and Trac-looping show high TSS signal enrichment (log2 fold change = 1.05 and 0.89, respectively, p < 2.2e-16), while Ocean-C data shows weaker enrichment (log2 fold change = 0.3, p < 2.2e-16). Given the low enrichment efficiency of Ocean-C, we did not include Ocean-C data in the following analysis. We also did similar analysis by comparing HiCAR data to the public HiChIP, PLAC-seq and DNase HiC data^12,13,21,24^. These comparisons showed that: (1) the signal enrichment of HiChIP/PLAC-seq at regulatory sequences depends on ChIP-antibody. For example, the antibody recognizing promoter mark H3K4me3, but not enhancer mark H3K4me1, shows TSS signal enrichment; and (2) as described in the original publication^24^, DNase HiC indeed showed no enrichment on the open chromatin regions. Thus, our results showed both HiCAR and Trac-looping can effectively enrich open chromatin anchored PETs, while HiChIP and PLAC-seq only captures a subset of cis-regulatory chromatin contacts. We further compared HiCAR data quality to the public Trac-looping^15^ and the in situ HiC data generated by 4DN consortium^25^ that were sequenced at similar depth (Fig 1D). Notably, HiCAR protocol requires much less input cells than Trac-looping and HiC, while producing 4.15-fold more (55.6% versus 13.4%) unique PET than Trac-looping. More importantly, compared to Trac-looping, HiCAR captured about 17-fold (18.3% versus 1.1%) more long-range (> 20kb) cis-PET that are the informative reads to identify long-range chromatin interactions (Fig 1E). Additionally, we evaluated the efficiency of HiCAR on capturing distal chromatin interaction across linear genomic distances at different scales. By examining genome wide average contract frequency, we found that HiCAR and *in situ* HiC show similar decay rate in capturing interactions with increased 1D distance, while Trac-looping captures more short-rage chromatin contract with in 7 kb but fewer chromatin interactions at long distance (Fig 1F). Overall, HiCAR outperforms existing methods to efficiently capture cis-regulagory chromaint contacts using low-input material and independent of antibody pulldown.

### HiCAR faithfully recapitulates the key features of 3D genome organization

Next, we asked if HiCAR data faithfully recapitulate the important features of genome structure. The 4DN consortium has generated deeply sequenced (total 6.2 billion raw reads) *in situ* HiC data using H1 hESC cells^21^. We took advantage of this 4DN *in situ* HiC dataset as a gold standard in the following analysis. First, we sought to determine if HiCAR could enrich the long cis-PET anchored on cis-regulatory elements. As shown in Fig 1G, we aggregated the HiCAR and *in situ* HiC chromatin contact matrix centered on the H1 hESC ATAC-seq peaks, CTCF binding sites, and peaks marked by H3K27ac, H3K4me1, H3K4me3, and H3K27me3. Within the 500kb window, HiCAR PET showed a clear stripe pattern extending from the aggregated peak centers. These results confirmed that HiCAR is an effective method to enrich pairwise long-range PET anchored on cREs. Second, we visually examined the chromatin contact heatmaps of HiC and HiCAR at different scales. As shown in Fig 1H, HiCAR generated a contact matrix highly similar to that of *in situ* HiC across different scales from chromosome, compartments, topological associated domains (TADs), down to the significant chromatin interactions at 5kb bin resolution. Notably, the HiCAR contact matrix, which was built from 488 million unique PET, revealed as much, if not greater, details on chromatin interactions compared to the deeply sequenced (2.53 billion unique PET) *in situ* HiC data. We observed similar compartment A/B segmentation, TAD boundaries, and significant chromatin interactions between the two datasets. At the genome-wide scale, the A/B compartment score, insulation score, and directionality index that are calculated from HiCAR and *in situ* HiC are well correlated with each other (Fig 1I). We concluded that HiCAR recapitulates the key features of high-order chromatin organization and effectively enriches the long-range chromatin interaction signals on the open chromatin sequences.

### HiCAR yields high-quality chromatin accessibility and transcriptome data with the same low-input biological sample

In a HiCAR experiment, the R2 reads of HiCAR DNA library are derived from the sequences targeted by Tn5 tagmentation, we speculate that the R2 reads can be treated as a single-end ATAC-seq reads to define the open chromatin regions across the genome. To test this, we took all the R2 reads from long range (>20kb) and the inter-chromosome trans-PET for 1D open chromatin analysis. The R2 reads from the short-range cis-PET were discarded in this analysis, because we found that these reads were biased towards the CviQI cutting site. Meanwhile, both the cytoplasm and nuclei RNAs from the same input cells were collected for RNA-seq analysis (Fig 1A). We made the RNA-seq library using a protocol modified from SMART-seq2^20^ (detailed in material and methods). After deep sequencing, we found that the HiCAR RNA-seq data and the filtered R2 reads were highly reproducible between biological replicates (Fig S1C). We further compared the HiCAR RNA-seq and the 1D open chromatin signal to the public H1 hESC RNA-seq and ATAC-seq data generated by ENCODE^26^ and the 4DN consortium^25^. As shown in Fig 1J, we observed very similar patterns of signal on the genome browser by visualization. At the genome-wide scale, the HiCAR RNA-seq and 1D peak reads are highly correlated with bulk RNA-seq and ATAC-seq dataset (Fig 1K, 1L). We used MACS2^27^ to call 1D open chromatin peaks using HiCAR R2 reads. As shown in Fig 1M, we found 57,069 overlapped open chromatin peaks between ATAC-seq and HiCAR 1D peaks. Meanwhile, we also observed unique peak sets that were identified in ATAC-seq and HiCAR 1D peaks, respectively. Next, we compared the MACS2 P-value of the overlapping and unique open chromatin peaks identified by ATAC-seq and HiCAR R2 reads. We found that the overlap peaks are associated with more significant P-value (MACS2) in both ATAC-seq and HiCAR 1D peak datasets (Fig 1N), suggesting HiCAR 1D peak analysis captures the high-confident peaks identified by ATAC-seq. Furthermore, we ranked the ATAC-seq peaks based on their P-value, and found that HiCAR 1D peaks covered more than 82% of high-confident ATAC-seq peaks with −log10 (P-value) > 8 (Fig S1D). Taken together, these results show that HiCAR can yield high-quality transcriptome and chromatin accessibility data using the same batch of low-input (100,000) crosslinked cells.

### HiCAR is a robust and sensitive method to identify cis-regulatory significant chromatin interactions

Our motivation for developing HiCAR is to identify the comprehensive long-range significant chromatin interactions anchored on cREs at high-resolution. We applied MAPS, a statistical and computational method recently developed for HiChIP/PLAC-seq loop call^28^, to our HiCAR datasets generated using H1 hESC. With MAPS, we first removed the potential systemic biases from the contact matrix, including GC content, mappability, 1D chromatin accessibility, and restriction enzyme cutting site. Finally, we identified 46,792 significant (FDR < 0.01) significant chromatin interactions at 5kb resolution.

After identifying these cis-regulatory loops, we first sought to estimate the true positive rate (sensitivity) of HiCAR in detecting known chromatin interactions. There is no “gold standard” set of true positive loops, so we decided to compare HiCAR loops to chromatin loops identified by well-established methods such as in situ HiC, PLAC-seq/HiChIP in matched cell types. Fortunately, there is high-quality public in situ HiC and H3K4m3 PLAC-seq data from H1 hESCs, as well as the CTCF HiChIP data generated in a recent study using H9 hESC^21,29^. Due to the lower sequencing depth of some of these public datasets, we decided to compare chromatin loops at 10kb resolution instead of 5kb resolution. In situ HiC and HiChIP/PLAC-seq data were processed using 10kb bins by HICCUPS^30^ and MAPS^28^, respectively. As shown in Fig 2A, visual examination of loop calls indicated that HiCAR loops showed striking similarity to loops identified by these well-established and widely used methods. Importantly, it appeared that in situ HiC, H3K4me3 PLAC-seq and CTCF HiChIP represent only a subset of the loops identified by HiCAR. To further quantify the sensitivity of HiCAR loops, we filtered the loops identified from in situ Hi-C, HiChIP/PLAC-seq, and only kept the loops with at least one anchor associated with an ATAC-seq peak for the following analysis. These loops were defined as the “testable loops” that we could compare to HiCAR loops (Fig 2B). We found HiCAR loops identified 92%, 81% and 69% of the “testable loops” identified by in situ HiC, H3K4me3 PLAC-seq, and CTCF HiChIP, respectively. These results suggest HiCAR is a highly sensitive method in detecting “known” chromatin loops identified by other well-established methods.

**Figure 2.**
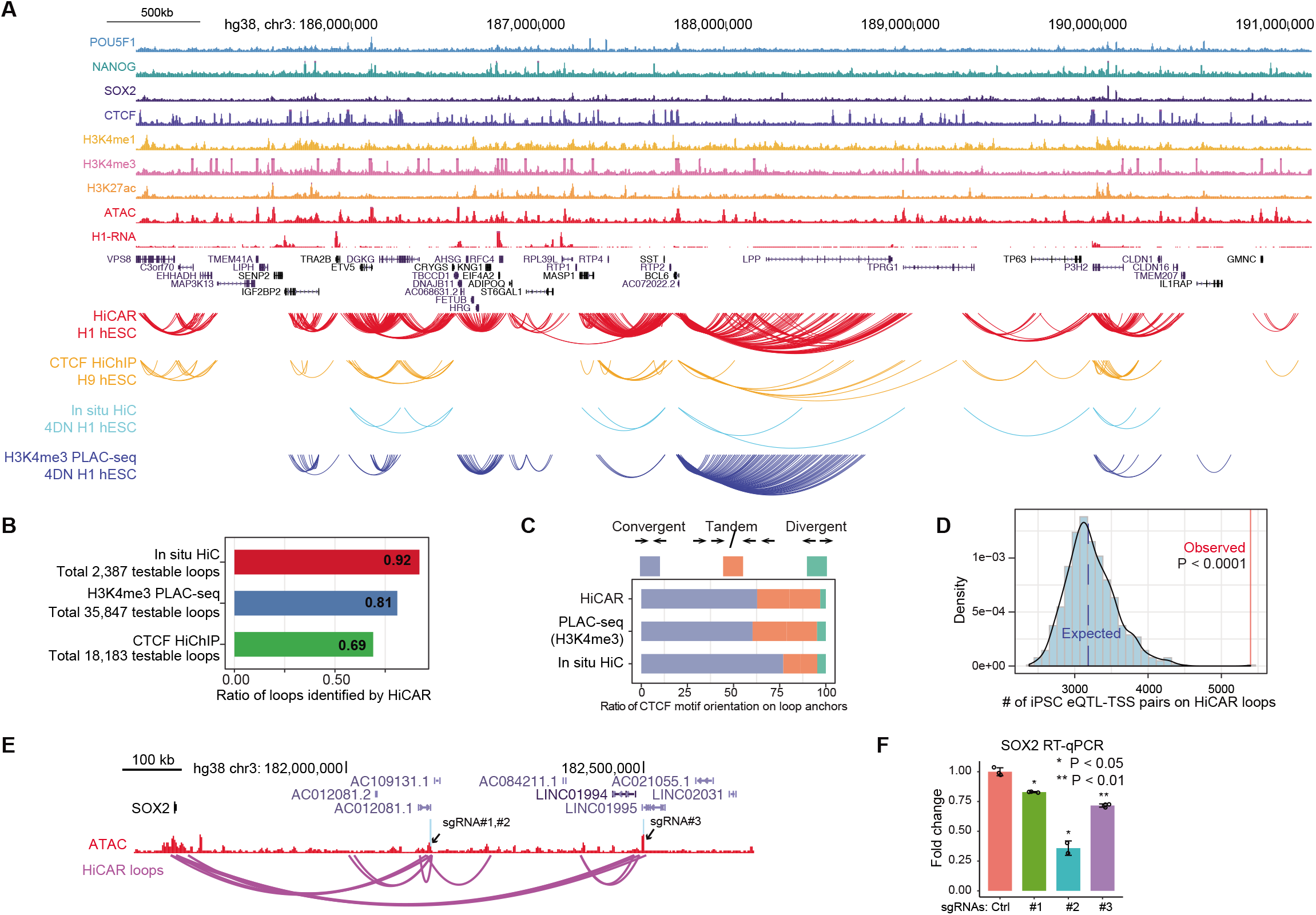
HiCAR recapitulates the spatial cREs interaction landscape. **(A)** UCSC Genome Browser visualization of NANOG, SOX2, CTCF, H3K4me1, H3K4me3 chip-seq, polyA RNA-seq, ATAC-seq and interaction/interactions identified from HiCAR, PLAC-seq-H3k4me3 and *in situ* HiC in H1 ESC cells and HiChip-CTCF in H9 ESC cells. **(B)** Percentage of detectable interactions in in-situ HiC, PALC-seq-H3K4me3 and HiChip-CTCF recovered by HiCAR datasets. **(C)** The ratio of convergent, tandem and divergent CTCF motif orientation at HiCAR, PALC-seq-H3K4me3 and *in situ* HiC interaction anchors. **(D)** Enrichment of eQTL-association in HiCAR loop regions. The red line represents the observed number of eQTL associations matched to HiCAR interactions. The histogram represents the distribution matched eQTL association to randomly sampled distance matched DNA regions (n = 10000). (one-sided empirical p-value < 0.0001). **(E)** UCSC genome browser shows the ATAC-seq and HiCAR identified interactions around SOX2 gene locus. Designed CRISPR-dCas9-KARB sgRNAs targeting region is highlighted by lightblue. **(F)** Barplot shows the SOX2 expression level normalized to GAPDH in the control and three sgRNA targeted samples. Data show represent three independently repeated experiments.

Next, we tried to assess the true positive rate for HiCAR loops. However, due to the lack of a complete list of true interactions in H1 hESC, we instead asked whether HiCAR loops recapitulate the known feature of chromatin interactions. Based on the loop exclusion model, it is known that CTCF/Cohesin-associated loops have a preference for convergent CTCF motif orientations at loop anchors^9^. Thus, we checked the CTCF motif orientation of the testable MAPS-identified interactions from HiCAR, and found the convergent CTCF motif rate is 62.8%, similar to the ratio of convergent CTCF motif observed in PLAC-seq (60.3%), but lower than in situ HiC loops (76.9%) (Fig 2C). We reasoned that such difference could be due the fact that HICCUPS uses local background for loop call, and only identifies the most significant loops. Furthermore, we explored the functional relationship of chromatin loops identified by HiCAR by examining whether HiCAR loops are enriched for expression quantitative trait loci (eQTL) and their associated gene/TSS identified in human pluripotent stem cells^31^. Indeed, we observed 5368 eQTL-TSS pairs overlapped with HiCAR loops, whereas 3228 eQTL-TSS pairs were expected to overlap with randomly selected pairwise genomics regions with matched distance to HiCAR loops (Fig 2D, empirical p value < 0.0001, detailed in Material and Methods). The significantly enriched eQTL-TSS pairs at HiCAR loops provided additional evidence suggesting the functional role of HiCAR loops in regulating gene expression. Finally, to directly test the causal role of HiCAR interactions, we selected three predicted SOX2 enhancers that form long-range interactions (Fig 2E, two enhancers at 430 kb, one enhancer at 788 kb) with the SOX2 promoter, and designed sgRNAs to specifically direct the epigenetic silencer dCas9-KRAB to the three candidate SOX2 enhancers. As shown in Fig 2F, epigenetic silencing of all three SOX2 enhancers led to significant downregulation of SOX2 mRNA expression level, showing that HiCAR interactions indeed reflect a functional interaction between enhancer and gene promoters. Taken together, our results showed that HiCAR is a robust, sensitive, and accurate way to identify the significant chromatin interactions between cREs.

### The promoters and enhancers forming long-range chromatin interactions exhibit significant correlated H3K27me3 ChIP-seq signal

Using HiCAR, we are able to identify 46,792 significant chromatin interactions at 5-kb resolution anchored on distinct epigenome states from one single assay. These interactions provide an unprecedented opportunity allowing us to interpret the regulation of epigenome, 3D genome, and gene expression. In order to obtain the fine-scale epigenome features associated with significant chromatin interactions, we took advantage of the rich epigenome datasets of H1 hESC generated by ENCODE and Epigenome Roadmap projects^2,17,26^ and public datasets from Cistrome data portal^32^, including 25 histone ChIP-seq and 43 TF ChIP-seq datasets (Supplementary table 1), and computed their corresponding signals on the HiCAR loop anchors. We are particularly interested in the question of whether these enhancers and promoters, which are proximal to each other in 3D, are associated with correlated certain epigenome features. To probe this question, we first took all the 5kb loop anchors overlapped with TSS, and defined them as “promoter” anchors. For the rest of anchors, if they overlap with H1 hESC enhancers annotated by chromHMM^2^, we defined them as “enhancer” anchors. The other anchors were defined as “other” anchors. By doing this, the 46,792 HiCAR interactions can be classified as promoter-enhancer (P-E, 24.8%), promoter-promoter (P-P, 22.7%), enhancer-enhancer (E-E, 10.6%), promoter-other (P-O, 20.3%), enhancer-other (E-O, 15.4%), and other-other (O-O, 6.17%) interaction (Fig 3A). Interestingly, we found that the interactions with both anchors located on promoter and/or enhancer (P-P, P-E, E-E) are significantly shorter than the interactions with one “other” anchor (P-O, E-O) (P < 0.001), while the “other-other” interactions are significantly longer (median distance 210 kb) than all other interactions (Fig 3B) (P < 4.55e-14). This analysis shows that enhancers and promoters tend to form relatively shorter significant chromatin interactions compared to other interactions. We speculate that the relatively shorter distance between enhancer and promoters might be helpful to maintain a robust and effective gene expression control program.

**Figure 3.**
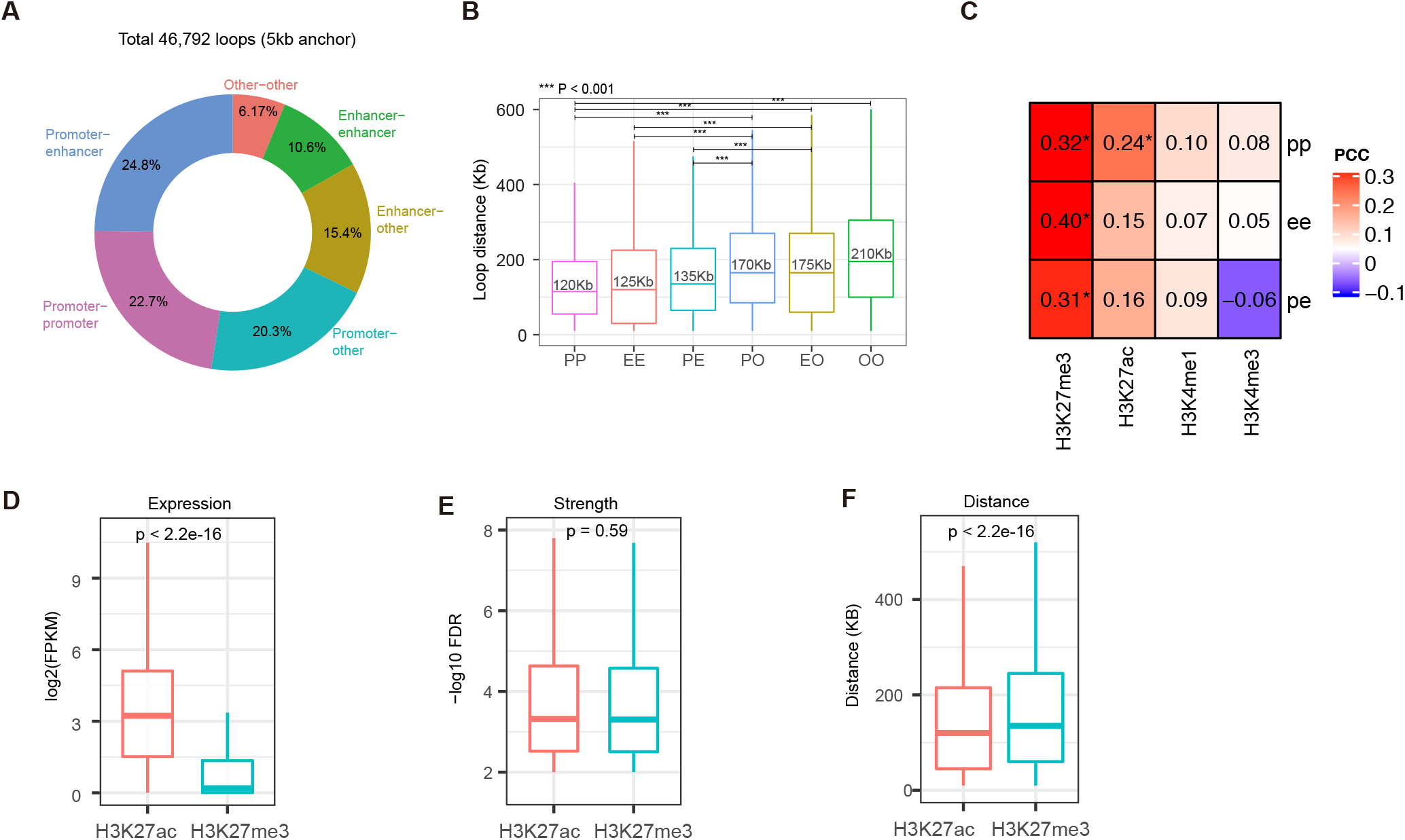
The posied cREs form extensive long-range chromatin interactions comparable to the active cREs. **(A).** The ratio of HiCAR chromatin interactions anchored on promoter, enhancer, and other cREs identified chromatin interac. **(B)** Boxplot shows the distribution of loop distances in each interaction group. **(C)** Heatmap shows the pearson correlation of indication histone marker signal between different groups of loop anchor regions identified by HiCAR. The significant correlation compared with shuffled background control are highlighted by “*”. **(D-F)** Boxplots show the gene expression level **(D)**, loop strength quantified by −log10 FDR **(E)**, and **(F)** loop distance between H3K27ac and H3K27me associated interactions.

The high-resolution HiCAR interactions allows us to test the hypothesis that the cREs which are linearly separated but positioned proximally in 3D space may also have coordinated status epigenetic modification. To test this hypothesis, we analyzed the histone modification signals on the anchor sequences of HiCAR interactions. Among these marks, H3K4me1 and H3K4me3 mark enhancer and promoter, respectively. We also examine the activation (H3K27me3) and repression/poised mark (H3K27me3). As shown in Fig 3C, we found that H3K27me3 showed a significant PCC correlation on promoter-promoter, promoter-enhancer and enhancer-enhancer interactions and H3K27ac showed a significant correlation on across P-P, P-E, and E-E anchors (PCC = 0.32, 0.31, and 0.40, P < 0.001). Interestingly, H3K27ac activation signal only shows significant correlation between P-P anchors (PCC = 0.24, P < 0.001), while the signal of H3K4me1 and H3K4me3 modification were not significantly correlated on these interactive loop anchors. This analysis suggested that long-range chromatin interaction may play a role in facilitating the H3K27me3 modification of the interacting enhancers and promoters.

### The poised/bivalent chromatin regions are associated with extensive interactive activity comparable to the activated chromatin state

H3K27me3 modification marks poised/bivalent chromatin state while H3K27ac is a widely used mark for active enhancer and promoter. Intrigued by the observation that H3K27me3 modifications are significantly correlated on the HiCAR interaction anchors, we decided to further examine the features of HiCAR interactions anchored on the poised cREs versus active cREs. We first identified 14,845 and 10,287 HiCAR interactions whose anchor sequences overlap with H1 hESC H3K27ac and H3K27me3 ChIP-seq peaks, respectively. Next, we identified the genes whose promoter sequences are located on these anchor sequences. As expected, genes associated with H3K27ac anchors are expressed at a much higher level compared to the genes expressed from H3K27me3-marked anchors (Fig 3D, p < 2.2e-16). Furthermore, we compared the loop strength quantified by −log10 FDR (output from MAPS) between two types of interactions. To our surprise, the HiCAR interactions associated with H3K27me3 anchors showed a similar distribution of FDR that are indistinguishable from those interactions anchored at active cREs marked by H3K27ac (Fig 3E, p = 0.59). Additionally, we compared the linear genomic distance of H3K27ac versus H3K27me3 anchored interactions. We found that the interactions with H3K27me3 modification are significantly longer (median distance 145 kb) than the H3K27ac interactions (median distance 125 kb) (Fig 3F, p < 2.2e-16). In order to gain further insight of the biological function involved in the two types of interactions, we performed Gene Ontology analysis. GO analysis showed that H3K27ac anchors are enriched for GO terms such as gene expression, metabolic, chromatin organization, and stem celll proliferation/maintenance, while the H3K27me3 anchor sequences are enriched for GO terms important for tissue and organ development. Our finding that the poised/H3K27me3 mark cREs are associated with extensive, long-range chromatin interaction is consistent with a recent study showing that H3K27me3-associated DNA interactions tend to span long-range and even across multiple topologically associated domains in mouse ESCs and human induced pluripotent stem cells^33^. However, these H3K27me3 anchored interactions were identified by H3K37me3 HiChIP analysis, thus, it remains challenging to compare these “poised” interactions to the chromatin interactions associated with different epigenetic modifications. Based on our analysis, we concluded that the poised cREs (marked H3K37me3) can form extensive and significant chromatin interactions that are comparable to the active cREs.

### Identification of super interactive regulatory (SINTER) loci enriched for housekeeping genes and developmentally regulated genes

It has been reported that a subset of gene promoters and super-enhancers are associated with high levels of chromatin interactivity known as super-interactive-promoter (SIP) and frequently interacting regions (FIREs)^34,35^, suggesting the cREs may be associated with unique 3D genome features. Thus, we decided to examine the spatial interactive activity of cREs. To tackle this problem, we ranked all the loop anchors according to their cumulative interaction scores (detailed in material and methods), and identified a subset of anchors (n = 2096) with unusually high degrees of chromatin interactivity, which we termed as <u>s</u>uper <u>i</u>nteractive <u>r</u>egulatory (SINTER) loci (Fig 4A). Through Gene Ontology enrichment analysis, we found that the genes located on the SINTER includes the house keeping genes (histone gene clusters), genes important for chromatin organization, stem cell proliferation, and many developmental genes and GO terms related to tissue and organ development (DLX3, SIX3, ELF3, POU3F1, MEIS1, and MYOD1) (Fig 4B, Supp Table 2). The presence of SINTER suggests that 3D chromatin organization plays an important role to ensure the proper regulation of these genes important for development and hESC lineage specification. Interestingly, for those genes located on SINTER, we found that they could be expressed at very low or high levels (green curve in Fig 4C), showing a similar pattern to the genes expressed from regulator HiCAR interaction anchors (pink curve in Fig 4C) and the genes not located on HiCAR loop anchors (Fig S2C, P = 0.96). Next, we asked if there are any features that specifically enriched on SINTER versus the regulator HiCAR anchors. As shown in Fig 4D, we found that H3K27ac, H3K27me3, SUZ12, and H3K4me1 signals are significantly enriched on SINTER versus regular loop anchors. The pluripotency marker POU5F1 and CTCF signals were also highly enriched on SINTER versus regular anchors. Interestingly, Cohesin (RAD21) signal is not enriched on the SINTERs compared to the regular anchors.

**Figure 4.**
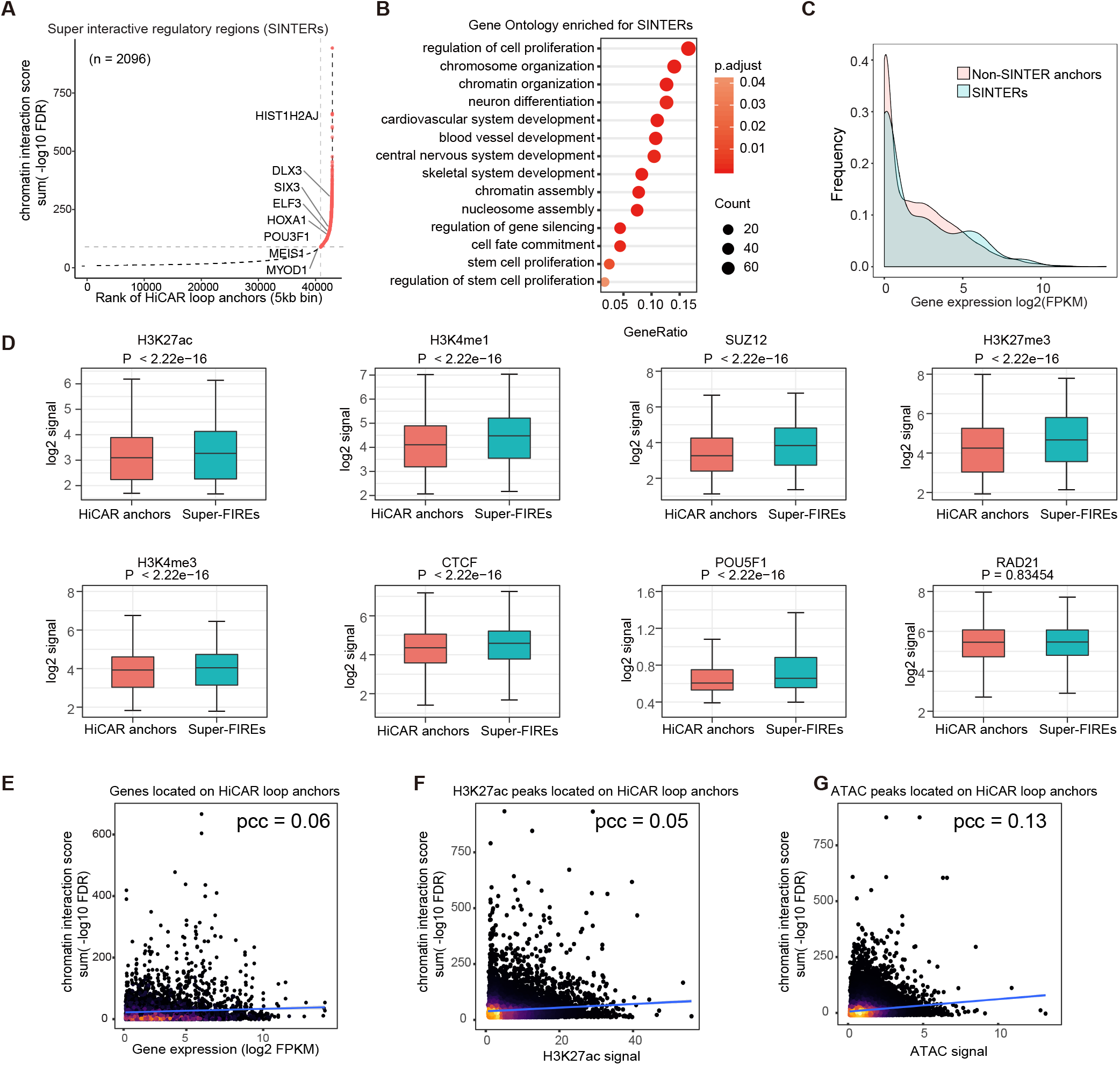
Analysis of cREs’ spatial interactive activity versus transcriptional activity. **(A)** Plot shows the ranked interaction score cumulative curve. The super interactive regulatory regions (SINTERs) are represented in red colour. **(B)** Gene Ontology enrichment analysis of SINTER loci associated genes. The dot represents the number of genes overlapped with each GO term and colour represents the degree of significant enrichment. **(C)** Density plot for the mRNA expression level of genes anchored SINTER loci (green) versus regulator HiCAR interaction anchors (pink). **(D)** Boxplot shows the indicated histone modification and TF binding at regular HiCAR interaction anchors (red) versus SINTER loci (green). **(E-G)** Scatter plots show the correlation of HiCAR anchors’ interactive score with the anchor associated gene expression level **(E)**, **(F)** active enhancer mark H3K27ac ChIP-seq peak average signal, and **(G)** ATAC-seq peak average signal.

Intrigued by the observation the highly interactive SINTER loci does not exhibit high level gene expression, we further asked, at the genome-wide scale, whether the cis-regulatory chroamtin loci’s spatial interaction activity correlate with their transcriptional activity. To probe this question, we first took the genes with promoters located on the HiCAR interaction anchors. We found that the mRNA levels of the genes expressed from HiCAR interaction anchors did not correlate with anchors’ spatial interactive activity (Fig 4E, PCC = 0.06). Further analysis showed that neither H3K27ac (Fig 4F) ChIP-seq signal or chromatin accessibility (Fig 4G) (ATAC-seq peak signal) were correlated with anchor’s spatial interactive activity (PCC = 0.05 and 0.13, respectively). Taken together, we identified 2096 SINTER enriched for housekeeping genes and developmentally regulated genes. Mechanistically, CTCF, but not Cohesin, may play a unique role to establish these SINTER with highly enriched signals of active and poised enhancers and promoters. Our analysis also revealed an interesting finding that the spatial activity of cRE does not correlate with its transcriptional activity, enhancer activity, or chromatin accessibility.

## Discussion

In summary, we applied HiCAR to H1 hESCs, and took advantage of the rich epigenome datasets generated from the same cell type by ENCODE, Epigenome Roadmap, and 4DN consortiums for integrative analysis. Consistent with recent studies showing that H3K27me3 and PRC2 are associated with extensive chromatin interactions anchored on the genes important for development^33,36^. In our study, we also found that the poised/bivalent promoter/enhancer marked by H3K27me3 and/or SUZ12 are associated with significant chromatin interactions near genes important for development. Compared to other studies using HiChIP and ChIA-PET to capture chromatin interactions only associated with H3K27me3 modification or PRC2 binding, we were able to identify the cis-regulatory chromatin interactions associated with multiple epigenetic marks from one single assay. This unique feature of HiCAR allows us to compare cis-regulatory chromatin contacts associted with a variety of different histone modifications and TFs. Our data showed that both the activated cREs and the poised/bivalent cREs are highly active in forming significant long-range significant chromatin interactions. Importantly, we found that the spatial interactive activity of cREs does not correlate with their transcriptional activity, enhancer activity, or chromatin accessibility. Additionally, we identified 2,096 SINTER loci associated with exceptionally high level chromatin interactive activity. SINTER loci are often near house keeping genes and developmental genes, and enriched for H3K27ac, H3K27me3, H3K4me1, POU5F1, CTCF but not Cohesin. Another interesting finding from our analysis is that the significant chromatin interactions between enhancers and/or promoters are significantly shorter than the chromatin interactions involving cREs that are not enhancer or promoter. We speculate that the relatively shorter range interactions might be more efficient to maintain a robust gene expression control mechanism.

Most importantly, we showed that HiCAR is a robust and sensitive method to study cis-regulatory chromatin contacts. First, HiCAR can efficiently enrich the long-range cis-PET so it requires much less sequencing depth than *in situ* HiC to identify high-resolution significant chromatin interactions. Second, compared to HiChIP/PLAC-seq, HiCAR does not rely on ChIP-grade antibodies to pull down the chromatin interaction associated with one specific protein or epigenetic modification. HiCAR enables comprehensive analysis of open chromatin-anchored interactions that are associated with a variety of epigenetic modification, while HiChIP and PLAC-seq only capture a subset of significant chromatin interactions identified by HiCAR analysis. Third, compared to the status quo technology known as Trac-looping, with similar sequencing depth, HiCAR generates ~17-fold more informative long-range cis-PET starting from 1000-fold less input cells. Lastly, HiCAR is also a robust multi-omics method that can be used to generate high-quality transcriptome and chromatin accessibility with the same biological samples used for 3D genome analysis. This unique feature makes HiCAR broadly applicable to low-input biological samples such as clinical samples or the primary cells purified from animal models. Our strategy of applying HiCAR in H1 hESC to identify 46,792 high-confident cRE-anchored significant chromatin interactions at kilobase resolution clearly demonstrated the technical advantage of our robust and sensitive multi-omic approach.

## ACKNOWLEDGEMENTS

We thank Dr. Kenneth Poss, Brigid Hogan and David Gorkin for critical reading of the manuscript. The funding of this work is supported by Duke Whitehead Scholar (to YD) and NIH U01 HL156064 (to YD). Y.X. is supported by a fellowship from Regeneration Next Initiative at Duke University.

## Supplementary Figure legend

**Supplementary Fig 1.**
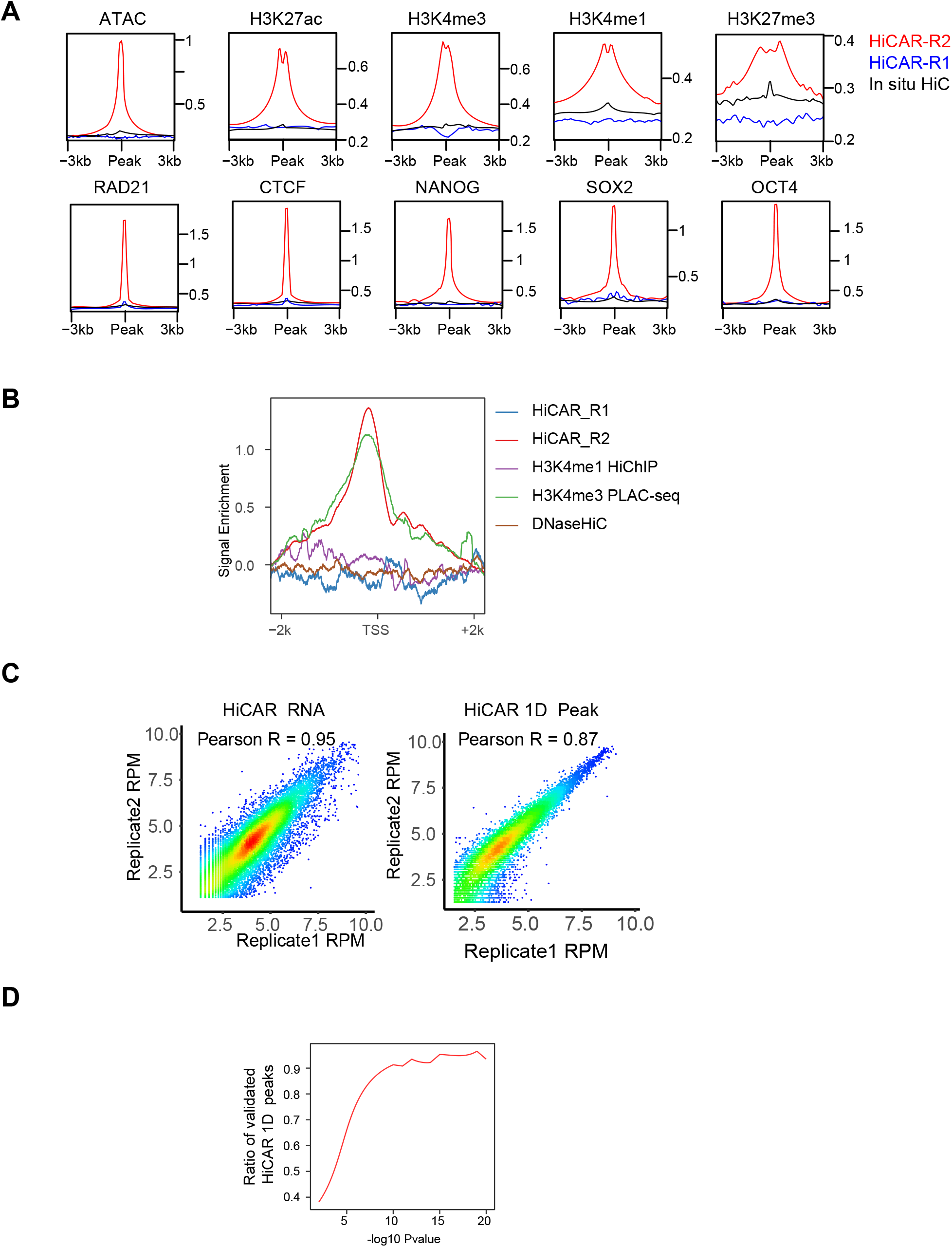
HiCAR signal enrichment and quality control. **(A)** The average signal of HiCAR R1 (Blue), R2 (Red) and *in situ* HiC (Black) around the indicated histone marker peak center region in 3kb windows. **(B)** Normalized signal in a window of 3kb around TSS sites for HiCAR R1 (blue), R2 (red), HiChIP H3K4me1 (purple), PLAC-seq H3K4me3 (green) and DNaseHiC (brown). **(C)** Scatter plots show the reads counts from two technical replicates of HiCAR RNA profile (left panel) and 1D Peaks (right panel). **(D)** Percentage of HiCAR ATAC peaks confirmed by bulk ATAC-seq dataset as a function of confidence scores.

**Supplementary Fig 2.**
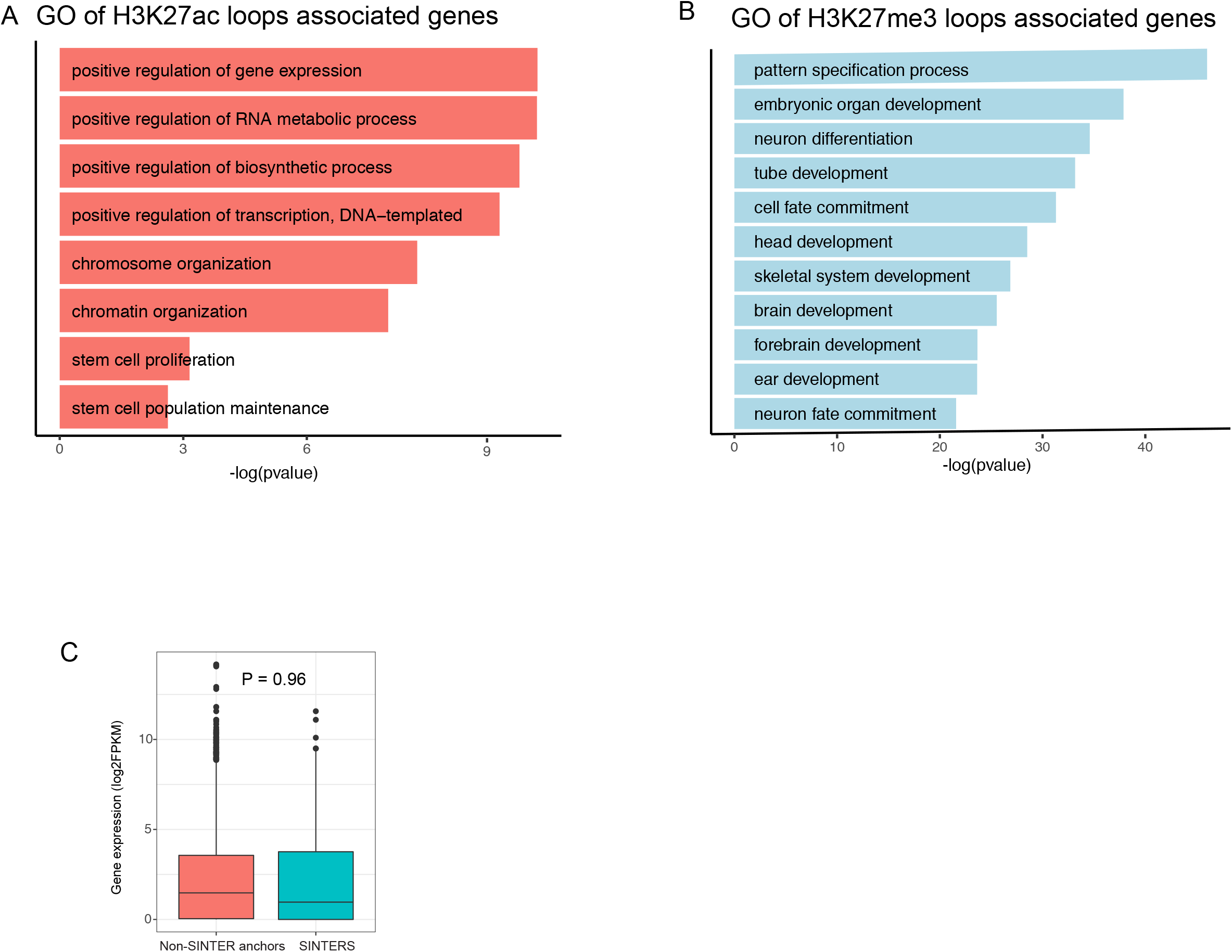
Correlation of interactive activity and gene expression. **(A-B)** Gene Ontology enrichment analysis of H3K27ac loops **(A)** and H3K27me3 loops **(B)** associated genes. **(C)** Boxplot shows the gene expression level between genes associated with SINTER anchor and non-SINTER HiCAR anchor.

## Method

### HiCAR data processing

HiCAR datasets were processed following the distiller pipeline (https://github.com/mirnylab/distiller-nf). Briefly, reads were aligned to hg38 reference genome using bwa mem with flags-SP. Alignments were parsed and paired end tags (PET) were generated using the pairtools (https://github.com/mirnylab/pairtools). PET with low mapping quality (MAPQ < 10) were filtered out. PET with the same coordinate on the genome or mapped to the same digestion fragment were removed. Uniquely mapped PET were then flipped as side1 with the lower genomic coordinate and aggregated into contact matrices in the cooler format using the cooler tools^37^ at delimited resolution (5kb, 10kb, 50kb, 100kb, 250kb, 500Kb, 1Mb, 25MB, 50MB,100MB). The dense matrix data were extracted from cooler files and visualized using HiGlass^38^. The R1 and R2 reads signal around TSS or peaks were calculated with EnrichedHeatmap^39^ before PET flipping.

### Compartments A and B, directionality and Insulation score

Compartmentalization, directionality index and insulation score was assessed using cooltools (https://github.com/mirnylab/cooltools). Briefly, eigenvector decomposition was performed on cis contact maps at 100-kb resolution. The first three eigenvectors and eigenvalues were calculated, and the eigenvector associated with the largest absolute eigenvalue was chosen. An identically binned track of GC content was used to orient the eigenvectors. The insulation score and directionality Index were computed by cooltools using ‘find_insulating_boundaries’ and ‘directionality’ function, respectively.

### Contact probability decaying curve

The curves of contact probability as a function of genomic separation were generated by pairsqc following the 4DN pipeline (https://github.com/4dn-dcic/pairsqc). Briefly, the genome is binned at log10 scale at interval of 0.1. For each bin, contact probability is computed as number of reads/number of possible reads/bin size.

### HiCAR RNA profile processing

Reads were aligned to hg38 with Hisat2^40^. Raw reads for each gene were quantified using featureCounts^41^. FPKM values were calculated for each gene.

### HiCAR 1D peak processing

Unique mapped HiCAR DNA library R2 reads were extracted before PET flipping. Only the long range (>20kb) and trans-PET were keeped for open chromatin analysis. MACS2^27^ were used to call ATAC peaks following the ENCODE pipeline (https://github.com/ENCODE-DCC/atac-seq-pipeline).

### CTCF motif orientation analysis

CTCF ChIP-seq peak list of H1 were downloaded from ENCODE and the searched for CTCF sequence motifs using gimme^42^ and CTCF motif (MA0139.1) from the JASPAR database^43^. We then selected a subset of interactions with both ends containing either a single CTCF motif or multiple CTCF motifs in the same direction. The frequency of all possible directionality of CTCF motif pairs, convergent, tandem and divergent, are then evaluated.

### Chromatin interaction calling

For HiCAR, PLAC-seq and HiChIP datasets, we used the MAPS^28^ to call the significant chromatin interaction from First, the interaction reads counts were extracted from cooler datasets at 5KB or 10Kb resolution. The interaction anchor bins were defined by the ATAC-signal peaks or corresponding ChIP-seq peaks called using MACS2^27^. MAPS applied a Poisson regression-based approach to normalize systematic biases from restriction sites, GC content, sequence mappability, and 1D signal enrichment. We grouped interactions that were located within 15 kb of each other at both ends into clusters and classified all other interactions as singletons. we retained only interactions with 6 or more reads and the significant interactions were defined by FDR < 0.01 for clusters and FDR < 0.0001 for singletons. For *in situ* HiC dataset, the .hic file is downloaded from 4DN data portal and HiCCUPS^30^ is applied to call interactions at 10Kb resolution with the following parameters: “-r 10000 -k KR -f .1,.1 -p 4,2 -i 7,5 -t 0.02,1.5,1.75,2 -d 20000,20000”.

### Loop classification

To dissect the epigenome properties of significant chromatin interactions, we first quantified the signal enrichment of 68 genomic datasets at each loop anchor (Supplementary Table 1). Loops with less than 10 epigenomes identified on their anchors are keeped for further analysis. We then built a covariance matrix for Principal Component Analysis (PCA) and dimension-reduced data from PCA were used to construct a KNN graph for loop clustering. This step is performed using the FindNeighbors function provided in the Seurat Package^44^. All loop clusters was visualized using UMAP projection.

### Gene Ontology enrichment analysis

We used Clusterprofile^45^ to examine whether particular gene sets were enriched in certain gene lists. GO categories with pvalue < 0.05 were considered significant.

### Cell culture and processing

WA01 were cultured in Matrigel (corning, 354230) coated plates with Stabilized feeder-free maintenance medium mTeSR™ Plus (STEMCELL, #05825). mTeSR™ Plus was changed every other day. For crosslinking, cells were washed once by PBS, then treated by accutase (biolegend, #423201) for 10mins at 37°C. After removing the accutase, cells were resuspended by DMEM. Formaldehyde was added to the final concentration of 1%, incubated at room temperature for 10mins. Glycine was added to the final concentration of 0.2M, incubated at room temperature for 10 mins to quench formaldehyde. Fixed cells were pelleted by centrifugation for 5 min at 4°C and washed with ice-cold PBS once

### Tn5 Purification

Briefly, Rosetta DE3 cells transformed with Tn5 expression plasmid pTXB1-Tn5 (Addgene #60240) were cultured in 500ml LB and incubated at 16°C overnight for protein induction. The bacteria were collected by centrifuge and resuspend by pre-cooled HEGX (40mM Hepes-KOH pH 7.2, 1.6M NaCl, 2 mM EDTA, 20% Glycerol, 0.4% Triton-X100, Roche Complete Protease Inhibitor), sonicated to release the protein. PEI (10% PEI, 4.44% HCl, 800mM NaCl, 20mM Hepes, 0.3mM EDTA, 0.2% Triton X-100, pH 7.2) were then added to the lysate in dropwise to precipitate the E. coli DNA. The lysate was then centrifuged and supernatant was loaded to Chitin column (BIO-RAD, #7372522). The column was rotated at 4°C for 2-3h then washed by HEGX buffer. 15ml HEGX buffer containing 100mM DTT was added to elute the protein. The column was incubated for another 24 hr at 4°C. The elution fraction was collected and concentrated to about 1ml by Amicon Ultracel 30K(Millipore, #UFC903024), then dialyzed twice by 1L dialysis buffer(100 HEPES-KOH pH 7.2, 0.2 M NaCl, 0.2 mM EDTA, 2 mM DTT, 0.2% Triton X-100, 20% glycerol) for 24h using dialysis membrane tube(Spectra, D1614-11). Then the protein was added 80% glycerol to a final concentration of 50%.

### Tn5 transposase assembly

To assemble Tn5, 50ul of 200mM ME-rev and 50ul of 200mM BfaI-truseqR1-pmeI-nextera7 were annealed by the following program: 95°C 5min, cool to 14°C with a slow ramp 1°C /min. The annealed adapter was mixed with Tn5 Transposase in 1: 1.5 molar ratio, the mixture was mixed by pipette and incubated at room temperature for 30mins.

### Cytoplasm and nuclei RNA libraries construction

The cytoplasm and nuclei supernatant was added 20% sds(final concentration 1%), mixed and incubated at 60°C for 30min. After adding 5M NaCl to final concentration 500mM, the sample was incubated at 68°C for at least 1.5h to reverse crosslink. After reverse crosslink, the RNA was purified by Phenol:Chloroform:Isoamyl Alcohol (25:24:1, v/v, SPECTRUM, #136112-00-0) extraction and 2.5X volume ethanol precipitation. The sample was dissolved by 21ul 10mM Tris-HCl (pH8.0). Then the sample was treated by 0,5ul DNaseI at 37°C for 30min to remove DNA in solution. Purified the RNA by 2X volume SPRI beads, dissolved RNA by 20ul 10mM Tris-HCl (pH8.0). Then take out 2.3ul RNA to make an RNAseq library using smartseq2 protocol^20^.

